# Deletion of M-opsin prevents “M cone” degeneration in a mouse model of Leber congenital amaurosis

**DOI:** 10.1101/803510

**Authors:** Hui Xu, Nduka Enemchukwu, Xiaoyue Zhong, Olivia Zhang, Yingbin Fu

## Abstract

Mutations in *RPE65* or lecithin-retinol acyltransferase (*LRAT*) disrupt 11-*cis*-retinal synthesis and cause Leber congenital amaurosis (LCA). In *Lrat*^−/−^ mouse model, mislocalized medium (M)-wavelength sensitive opsin was degraded whereas mislocalized short (S)-wavelength sensitive opsin accumulated before the onset of cone degeneration. The mechanism for the foveal medium (M)/long (L)-wavelength-sensitive cone degeneration in LCA is unknown. By crossing *Lrat*^−/−^ mice with a proteasome reporter mouse line, we showed that M-opsin enriched dorsal cones in *Lrat*^−/−^ mice exhibit proteasome stress due to the degradation of large amounts of M-opsin. Deletion of M-opsin relieves the proteasome stress and completely prevents “M cone” degeneration in *Lrat*^−/−^*Opn1sw*^−/−^ mice (a pure “M cone” LCA model, *Opn1sw*^−/−^ encoding S-opsin) for at least 12 months. Our results suggest that M-opsin degradation associated proteasome stress plays a major role in “M cone” degeneration in *Lrat*^−/−^ model. This finding may represent a general mechanism for “M cone” degeneration for multiple forms of cone degeneration due to M-opsin mislocalization and degradation. Our results have important implications for the current gene therapy strategy for LCA that emphasizes the need for a combinatorial therapy to both improve vision and slow photoreceptor degeneration.

## Introduction

Retinoid isomerase (RPE65) and lecithin-retinol acyltransferase (LRAT) are two key enzymes involved in the generation/recycling of 11-*cis*-retinal in the retinal pigment epithelium (RPE). Mutations in either gene lead to Leber congenital amaurosis (LCA), a severe inherited retinal degenerative disease characterized by severe loss of vision in childhood with early degeneration of foveal cones followed by rods (1, 2). Two mouse models *Rpe65*^−/−^ (LCA2) and *Lrat*^−/−^ (LCA14) mice, which have shown very similar phenotypes, and a *Rpe65*^−/−^ dog model have been widely used for mechanistic and therapeutic studies (3–11). These studies paved the way for the first successfully treated inherited retinopathy using gene therapy (12–14). However, follow-up studies found that photoreceptor degeneration continued even after the gene therapy intervention in LCA2 (11, 15–17), suggesting a need to understand the mechanism of photoreceptor degeneration to design improved treatment strategies.

In both LCA patients and animal models, both rod and cone functions are severely compromised due to a combination of 11-*cis*-retinal deficiency and photoreceptor degeneration. Early loss of foveal cones was reported in RPE65-deficient patients (1, 2). Despite extensive studies, the mechanisms underlying cone degeneration in LCA are still poorly understood. In mouse models, both medium-wavelength sensitive opsin (M-opsin) and short-wavelength sensitive opsin (S-opsin) fail to traffic from the cone inner segment to the outer segment and cone photoreceptors in the central/ventral retina degenerate rapidly (< 4 weeks) (5–7, 18). Dorsal cones degenerate slowly (> 6 months). We have shown previously that S-opsin was aggregation-prone in cell culture, and accumulated in cones leading to proapoptotic endoplasmic reticulum (ER) stress and rapid S-opsin enriched central/ventral cone (“S-cone”) degeneration in *Lrat*^−/−^ mice (8). Deletion of S-opsin reduced ER stress and completely prevented the rapid S-cone degeneration in *Lrat*^−/−^*S-opsin^−/−^* mice for at least 12 months (19). However, the mechanism for M-opsin enriched dorsal cone (“M cone”) degeneration is unclear. Understanding the mechanism for M/L (medium/long-wavelength) cone degeneration is important since the human fovea is dominated by both cone types, which are responsible for high-resolution daylight vision and color perception (20).

Our previous work has shown that mistargeted M-opsin was largely degraded whereas S-opsin was resistant to degradation in *Lrat*^−/−^ mice (8). M-opsin was shown to be degraded by the proteasome but not lysosome pathway in *Rpe65*^−/−^ mice (21). Degradation of large amounts of mistargeted M-opsin in cones will likely overload proteasome and cause proteasome stress. Interference with the proteasome inhibits both cytosolic protein degradation and ER-associated degradation (ERAD), which can lead to cell death. Indeed, the degradation of rod transduction proteins was shown to cause proteasome overload, ER stress, and rod degeneration (22–24). Moreover, decreased proteasomal activity causes rod degeneration (25), whereas increased proteasomal activity delays rod degeneration (26). However, the role of proteasome stress in cone degeneration has not been studied. Due to mixed expression of M- and S-opsins in mouse cones and the rapid degeneration caused by S-opsin in *Lrat*^−/−^ mice, we used the *Lrat*^−/−^*Opn1sw ^−/−^* mice as a pure “M cone” LCA model to study the role of M-opsin and proteasome stress in cone degeneration. We showed that deletion of M-opsin relieves the proteasome stress and prevents “M cone” degeneration in *Lrat*^−/−^*Opn1sw ^−/−^* mice for at least 12 months.

## Results and Discussion

Previous studies have shown that M-opsin was degraded in *Lrat*^−/−^ mice (8, 19) and that M-opsin degradation was mediated by the proteasome but not lysosome pathway (21). We hypothesize that the degradation of large amounts of mislocalized M-opsin can overload the ubiquitin-proteasome system and cause proteasome stress. To examine proteasomal insufficiency in *Lrat^−/−^* cones, we crossed *Lrat^−/−^* with a proteasome reporter mouse line Ub^G76V^-GFP to generate *Lrat^−/−^Ub^G76V^*-*GFP* mice. Accumulation of Ub^G76V^-GFP (a cytoplasmic substrate carrying a degradation signal) indicates proteasomal insufficiency (27). The accumulation of Ub^G76V^-GFP was detected by immunohistochemistry with an anti-GFP antibody (28). Cones from all regions of P18 *Lrat^−/−^Ub^G76V^*-*GFP* displayed robust reporter signal throughout cones, i.e., from outer segment to synaptic terminal in green [Cones were labeled with rhodamine-peanut agglutinin (PNA) in red] (Fig. 1). No cone specific-GFP signal was detected in control *Ub^G76V^*-*GFP*. Proteasomal insufficiency in central/ventral cones was likely caused by aggregated S-opsin mediated proteasome inhibition (29–32). Proteasome insufficiency in dorsal cones was likely caused by M-opsin degradation induced proteasome overload.

**Figure 1.**
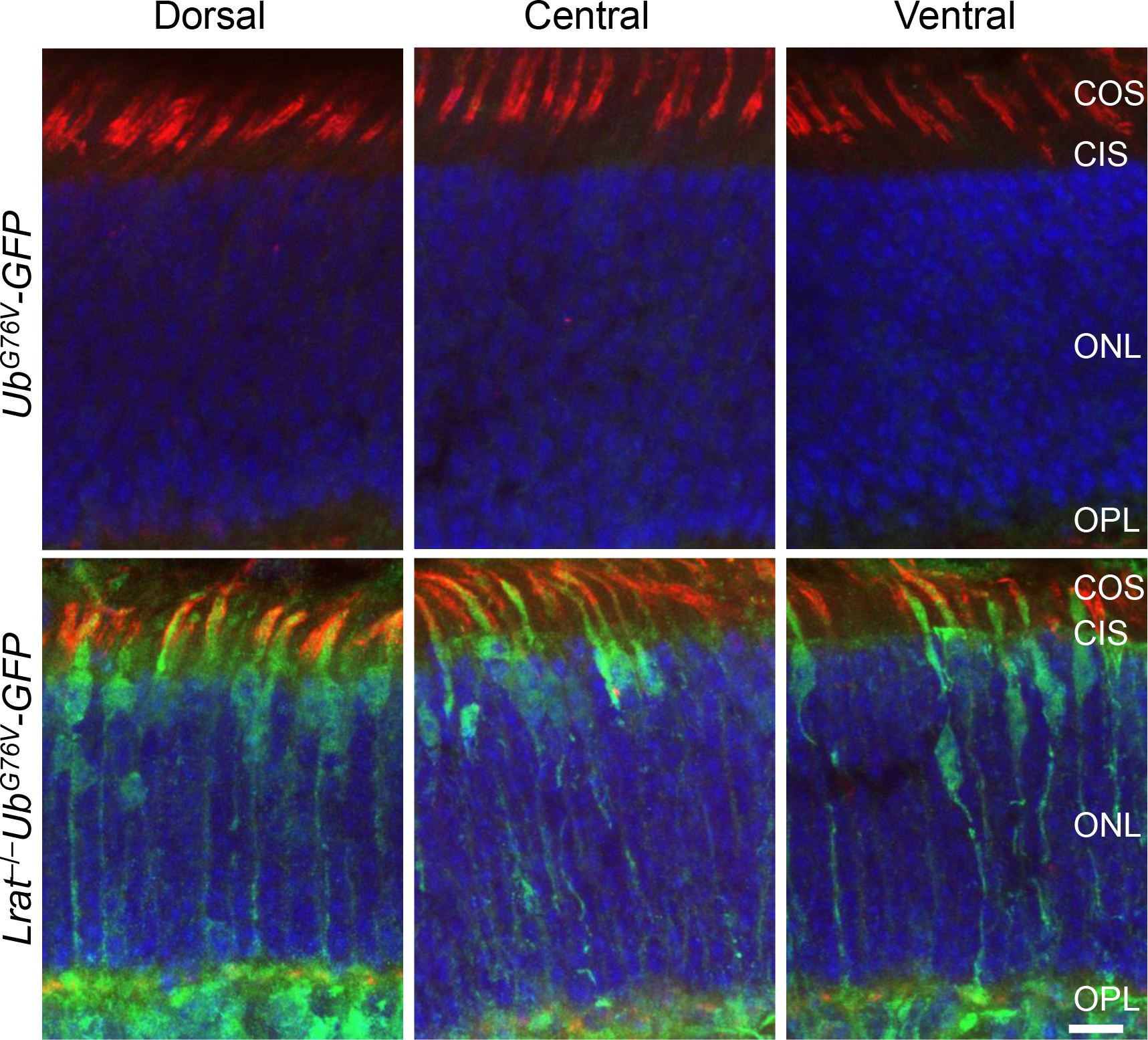
Cones in *Lrat*^−/−^ mice exhibit proteasome insufficiency. The retinal sections from P18 *Lrat^−/−^Ub^G76V^*-*GFP* and *Ub^G76V^*-*GFP* control mice were labeled with an anti-GFP antibody (in green). Cones were labeled with rhodamine-Peanut Agglutinin (in red). Nuclei were stained with 4’, 6-diamidino-2-phenylindole (DAPI). COS, cone outer segment. CIS, cone inner segment. ONL, outer nuclear layer. OPL, outer plexiform layer. Scale bar, 10μm.

To overcome the difficulties associated with the mixed expression of M and S-opsins in mouse cones and the rapid degeneration caused by S-opsin in *Lrat*^−/−^ mice, we generated the *Lrat*^−/−^*Opn1sw* ^−/−^ mice as a pure “M cone” LCA model to study the role of M-opsin in cone degeneration. We first generated *Opn1mw* ^−/−^ mice by CRISPR/Cas9. *Opn1mw* was disrupted by 1 nucleotide insertion in the second exon, which causes a frame shift and truncation at the 2nd exon (Fig. 2A &B, Supplementary Fig. 1). *Opn1mw* ^−/−^ mice (male *Opn1mw*^*Y*/-^ and female *Opn1mw* ^−/−^ mice) have normal body size and fertility. Immunohistochemistry and western blot confirmed the absence of M-opsin in the retina of *Opn1mw* ^−/−^ mice (Fig. 2 C & D). The expression level of S-opsin was increased in *Opn1mw* ^−/−^ mice (Figure 2C), which mirrors the result of M-opsin upregulation in *S-opsin*^−/−^ mice due to transcriptional compensation (33). We bred *Opn1mw* ^−/−^ with *Lrat^−/−^Opn1sw* ^−/−^ mice to generate *Lrat^−/−^Opn1sw ^−/−^Opn1mw* ^−/−^ mice. Dorsal cones of *Lrat^−/−^Opn1sw* ^−/−^ mice are morphologically unhealthy at 6 months with swollen outer and inner segment (Fig. 3A, white arrowheads). This became much worse at 12 months. Most remaining cones have lost their outer segment (Fig. 3A, white arrows) with broken or fragmented structure (Fig. 3A, red arrows). Compared with *WT*, *Lrat^−/−^Opn1sw* ^−/−^ mice have lost 26% (p<0.01) and 49% (p<0.001) cones at 6 and 12 months, respectively (Fig. 3B). In contrast, 97% and 94% dorsal cones survived in *Lrat^−/−^Opn1sw ^−/−^Opn1mw* ^−/−^ mice at 6 and 12 months, respectively, which is not significantly different from control *WT* and *Opn1sw* ^−/−^*Opn1mw* ^−/−^ mice, suggesting that M-opsin plays a major role in dorsal cone degeneration in *Lrat*^−/−^ mice. We have shown previously that deletion of S-opsin prevents ventral and central cone degeneration in *Lrat^−/−^Opn1sw* ^−/−^ mice up to 12 months. We confirmed this result in Fig. 2B. Previous work by others and us have shown cones can survive with little cone opsins (i.e. ventral cones in *Opn1sw* ^−/−^ and *Lrat^−/−^Opn1sw* ^−/−^ mice) (19, 33), our new data provide conclusive evidence that cones can survive with no cone opsins (i.e., 6- and 12-month *Opn1sw* ^−/−^*Opn1mw* ^−/−^ cones), which is quite different from rods that degenerate rapidly without rhodopsin (34).

**Figure 2.**
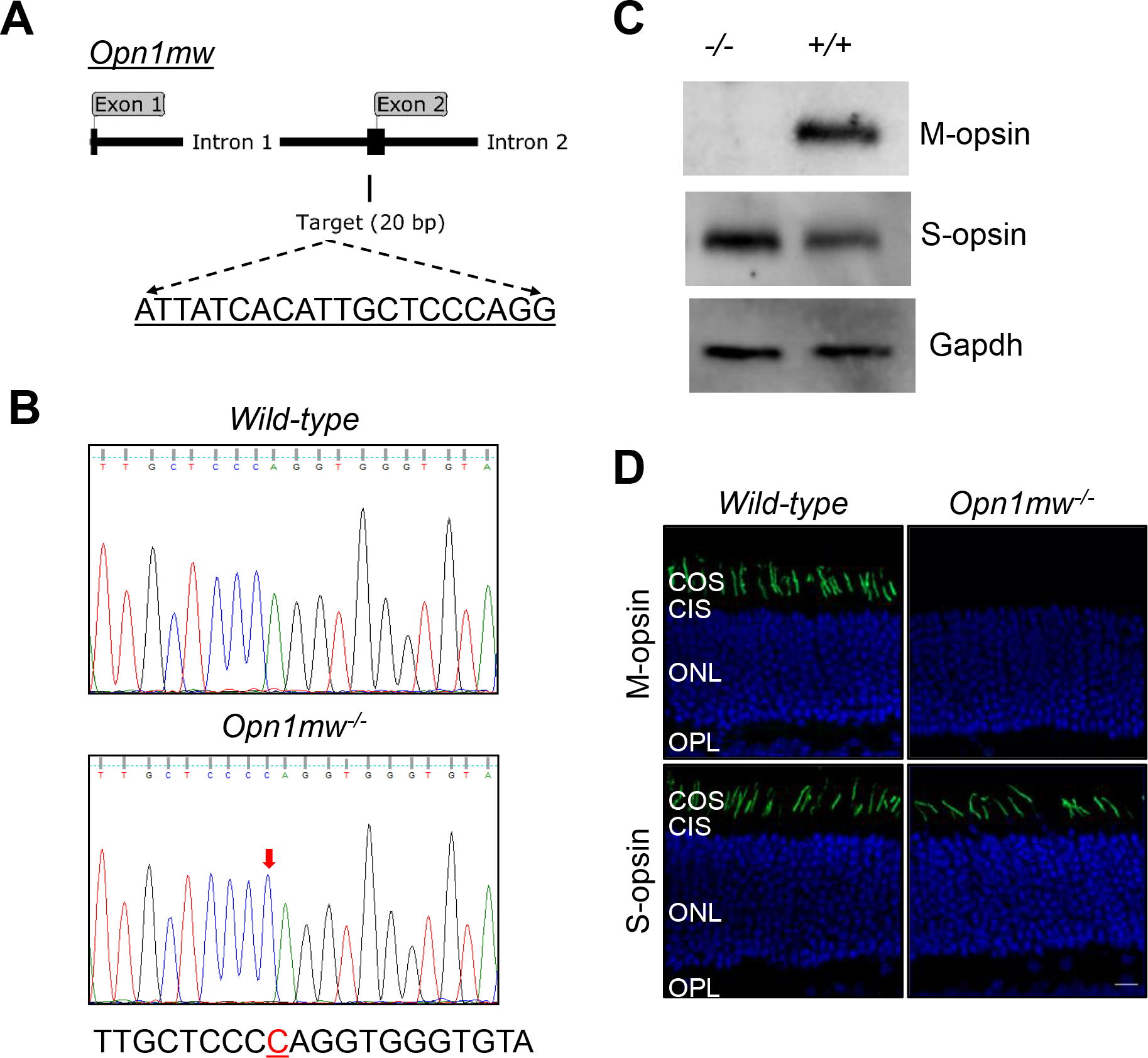
Generation and characterization of *Opn1mw* ^−/−^mice. **(A)** A schematic illustration on the location of the sgRNA targeting sequence in mouse *Opn1mw*gene. **(B)** DNA sequencing traces of *WT* and *Opn1mw* ^−/−^mice. There was a 1 bp insertion (red arrow) at the targeting site, which caused a frame shift and truncation (Supplementary Fig. 1). **(C)** Western blotting analysis of M-opsin and S-opsin from 1-month *WT* and *Opn1mw* ^−/−^retinas. Gapdh was used as a loading control. **(D)** Immunolabeling of M-opsin and S-opsin in the retinal sections of 1-month *WT* and *Opn1mw* ^−/−^mice. Nuclei were stained with DAPI (blue). Scale bar, 10 μm.

We bred *Lrat^−/−^Opn1sw^−/−^Opn1mw*^−/−^ and *Lrat^−/−^Opn1sw*^−/−^ mice with Ub^G76V^-GFP to generate *Lrat^−/−^Opn1sw^−/−^Opn1mw^−/−^Ub^G76V^*-*GFP* and *Lrat^−/−^Opn1sw^−/−^Ub^G76V^*-*GFP* mice to assess the role of proteasome stress in “M-cone” degeneration. In P18 *Lrat^−/−^Opn1sw*^−/−^ mice, dorsal cones displayed strong GFP signal indicating proteasome inefficiency (Fig. 3C, white arrows). In contrast, dorsal cones from P18 *Lrat^−/−^Opn1sw^−/−^Opn1mw^−/−^Ub^G76V^*-*GFP* mice exhibited markedly decreased GFP signal, suggesting M-opsin degradation plays a major role for proteasome stress in M cones. Interestingly, GFP signal in the ventral and central cones in *Lrat^−/−^Opn1sw^−/−^Opn1mw^−/−^Ub^G76V^*-*GFP* mice was also decreased compared with those of *Lrat^−/−^Opn1sw*^−/−^ mice (Supplementary Fig. 2). One explanation is that in the absence of S-opsin, M-opsin is increased due to transcriptional compensation and causes proteasome stress in central and ventral cones (Fig. 2C) (33). The residual GFP signal in *Lrat^−/−^Opn1sw^−/−^Opn1mw^−/−^Ub^G76V^*-*GFP* mice compared with *WT* mice is likely caused by proteasome stress associated with the degradation of other cone phototransduction proteins (see below).

**Figure 3.**
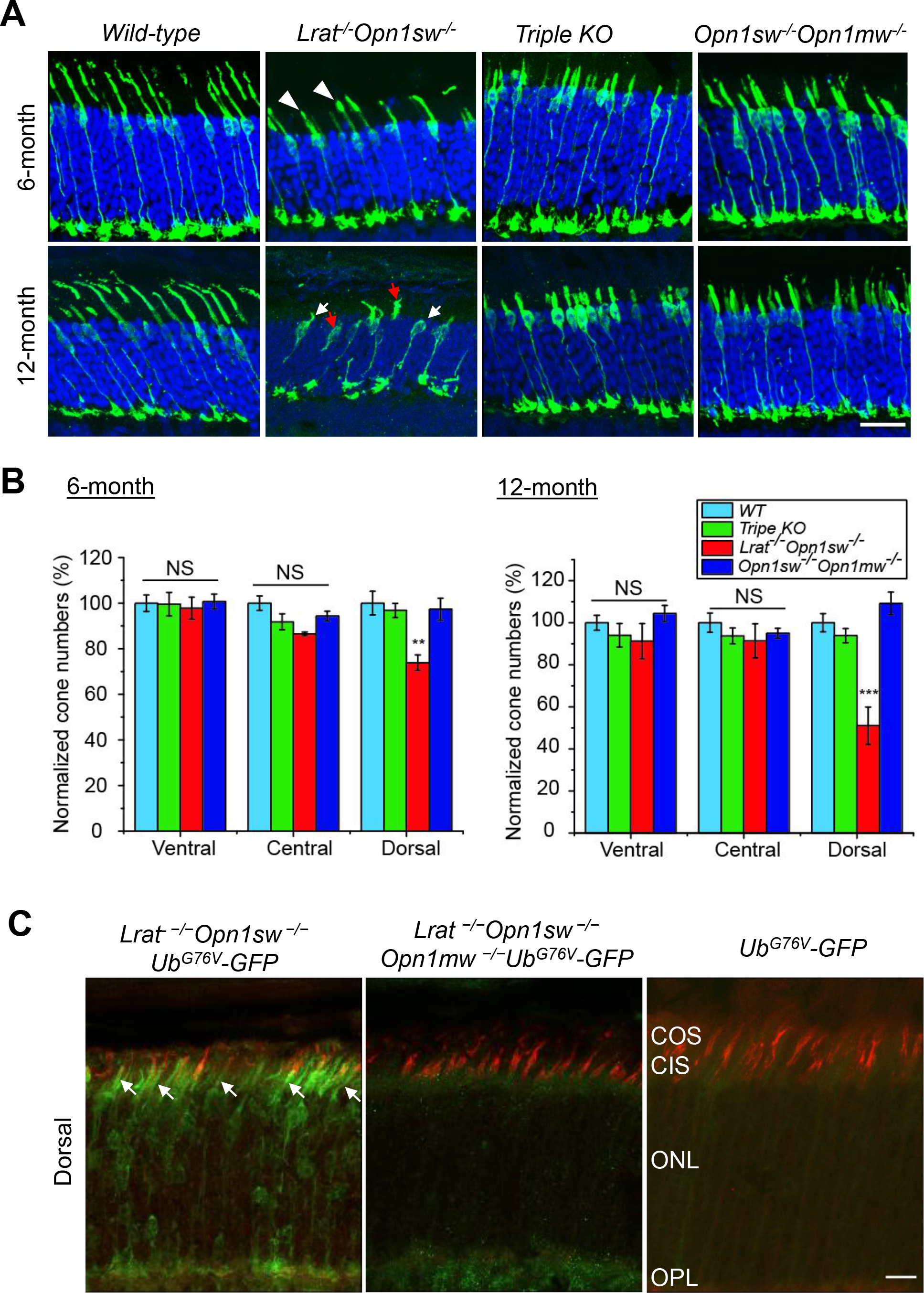
M-opsin deletion prevents dorsal “M cone” degeneration in *Lrat*^−/−^*Opn1sw*^−/−^*Opn1mw*^−/−^ mice. **(A)** Retinal sections from 6 months and 12 months old *WT*, *Lrat*^−/−^*Opn1sw*^−/−^, *Lrat*^−/−^*Opn1sw*^−/−^*Opn1mw*^−/−^(*Triple KO*), *and Opn1sw*^−/−^*Opn1mw*^−/−^ mice were labeled with anti-mouse cone arrestin antibody (in green). Representative retinal images from the dorsal regions of indicated genotype were shown. Nuclei were stained with DAPI. Scale bar, 20 μm. **(B)** Quantitative results of relative cone numbers at the ventral, central and dorsal retina from the indicated genotype. Data (mean ± SEM) were normalized to WT dorsal region. For 6-month data, N= 4 for *WT*, *Lrat*^−/−^*Opn1sw*^−/−^, *and Opn1sw*^−/−^*Opn1mw*^−/−^. N= 7 for *Triple KO*. For 12-month data, N=6 for *WT* and *Triple KO* for ventral WT (N=5); N= 4 for *Lrat*^−/−^*Opn1sw*^−/−^, and *Opn1sw*^−/−^*Opn1mw*^−/−^ except for central *Lrat*^−/−^*Opn1sw*^−/−^ (N=5). ** p<0.01, *** p<0.001 by one-way ANOVA with Tukey post hoc analysis, in which the dorsal cone numbers of *Lrat*^−/−^*Opn1sw*^−/−^ mice were significantly decreased compared with all other three genotypes at both 6-month and 12-month. NS, not significant. **(C)** M-opsin deletion reduces proteasomal stress in dorsal “M cones” in *Lrat*^−/−^*Opn1sw*^−/−^ mice. The reporter Ub^G76V^-GFP signal from the dorsal retinas of P18 *Lrat*^−/−^*Opn1sw*^−/−^ *Ub^G76V^*-*GFP* mice, *Lrat*^−/−^*Opn1sw*^−/−^*Opn1mw*^−/−^*Ub^G76V^*-*GFP*, and *Ub^G76V^*-*GFP* control mice were labeled with an anti-GFP antibody (in green). Cones were labeled with rhodamine-PNA (in red). Scale bar, 20 μm in A and 10 μm in C.

Several membrane-associated proteins, e.g., M/S opsins, G protein-coupled receptor kinase 1 (GRK1), cone transducin alpha subunit (G_α_t2), involved in cone phototransduction fail to traffic to the outer segment of *Lrat*^−/−^ or *Rpe65*^−/−^ cones properly (7). The mistrafficked proteins are degraded through a posttranslational mechanism (7), except that S-opsin aggregates and resists proteasome degradation (8). We examined the subcellular localization of GRK1 and G_α_t2 in the retinas of *Lrat^−/−^Opn1sw^−/−^Opn1mw*^−/−^, *Opn1sw*^−/−^*Opn1mw*^−/−^, *Lrat^−/−^Opn1sw*^−/−^, and *WT* mice. GRK1 was expressed in both rods and cones (Supplementary Fig. 3, top left panel, white arrows indicating cone signal) in *WT*. In sharp contrast, no GRK1 signal was detected in cones in *Lrat^−/−^Opn1sw^−/−^Opn1mw*^−/−^, *Opn1sw*^−/−^*Opn1mw*^−/−^, and *Lrat^−/−^Opn1sw*^−/−^ mice (Supplementary Fig. 3, top right three panels). G_α_ t2 was markedly reduced in *Lrat^−/−^Opn1sw*^−/−^ mice (Supplementary Fig. 3, 2nd bottom left panel) as we have shown previously (8), and was undetectable in *Lrat^−/−^Opn1sw^−/−^Opn1mw*^−/−^ and *Opn1sw*^−/−^*Opn1mw*^−/−^ mice (Supplementary Fig. 3, bottom right two panels) compared to the robust signal in *WT* cones (Supplementary Fig. 3, bottom left panel, red arrows). These results suggest that the degradation of many cone membrane-associated proteins other than M-opsin does not play a significant role in cone viability likely due to their much smaller quantities than M-opsin although it may still cause some degree of proteasome stress (see Supplementary Fig. 1).

The major finding of this study is that deletion of M-opsin relieved proteasome stress and prevented the dorsal “M cone” degeneration in *Lrat^−/−^S-opsin*^−/−^ mice for at least one year. In the absence of 11-*cis*-retinal, daily degradation of large amounts of M-opsin could overload the proteasome and cause chronic proteasomal insufficiency, which may lead to the accumulation of misfolded proteins in the ER by inhibiting ERAD and triggering chronic ER stress in M cones. On the other hand, chronic proteasomal insufficiency may also affect normal cellular function (35, 36). Indeed, proteasome overload or impairment has been implicated in several types of rod degeneration associated with protein misfolding or mistargeting (23, 37–39). To our knowledge, this is the first study to examine the role of proteasome stress in cone degeneration. Since a number of mouse models of cone dystrophy with cone opsin mislocalization exhibited M-opsin degradation and slower dorsal cone degeneration, e.g., cone cyclic nucleotide-gated channel A subunit (*Cnga3*) knockout (40), GC1 (*Gucy2e*) knockout (LCA1) (41, 42), Rd 3 (LCA12) (43), AIPL1 knockout (*LCA4*) (44), and kinesin 3A (*Kif3a*) knockout in cones (45), M-opsin degradation associated proteasome stress may represent a general mechanism for dorsal cone degeneration in these models (Fig. 4).

**Figure 4.**
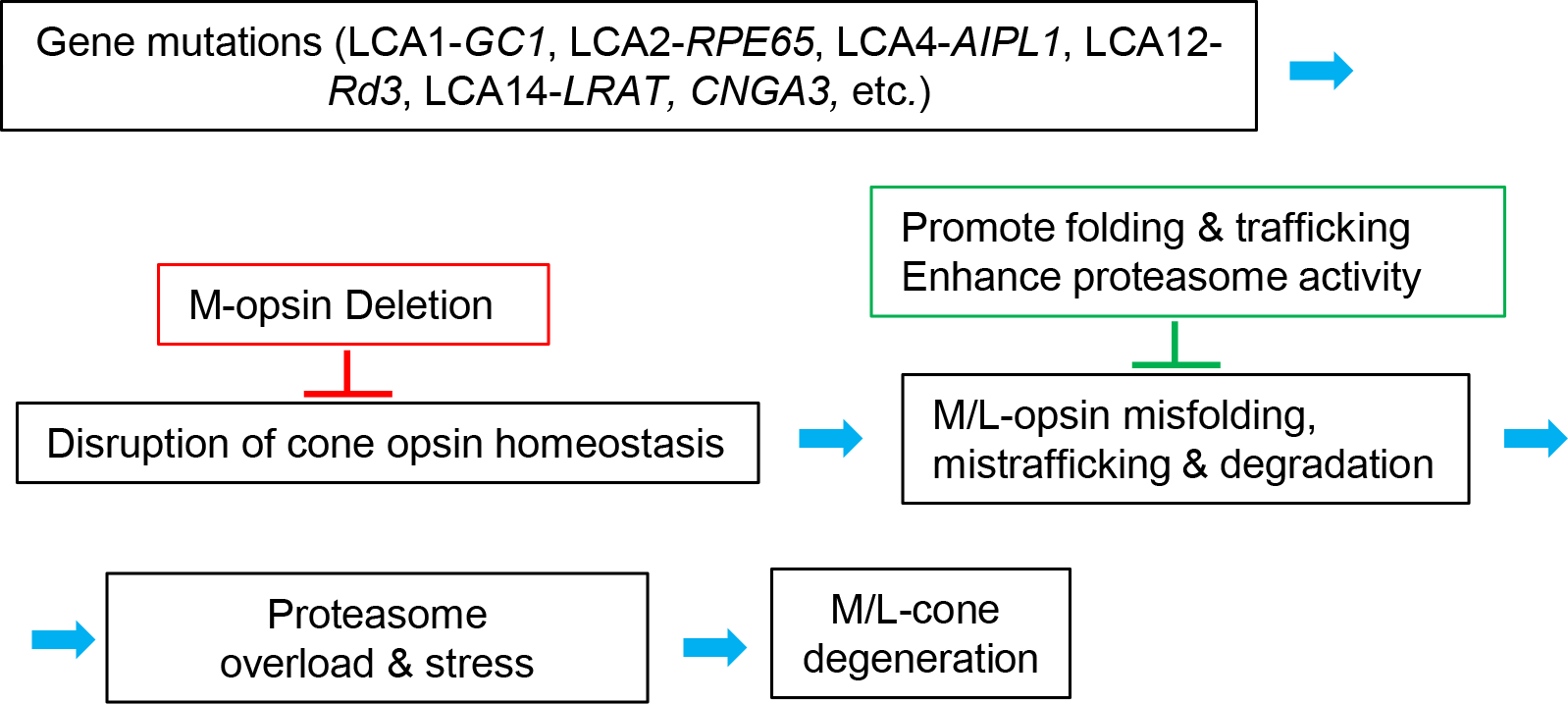
A schematic on the role of M-opsin in multiple forms of cone degeneration. Mislocalized M/L-opsins are degraded, which causes proteasome stress and dorsal “M cone” degeneration. Deletion of M-opsin prevents dorsal “M cone” degeneration. One strategy to protect M/L-cones from degeneration is to promote the folding and targeting of M/L-opsins and to enhance proteasome activity.

The most promising treatment for LRAT *or* RPE65 LCA is to use gene augmentation therapy with a normal copy of *LRAT* (or *RPE65*) to restore the visual cycle. In 2017, Luxturna was approved by Food and Drug Administration (FDA) for the treatment of RPE65-LCA (LCA2), making it the first directly administered gene therapy approved in the United States that targets a genetic disease caused by mutations in a specific gene. However, there were reports that photoreceptors continued to degenerate and vision benefit faded within 3 years despite initial vision gain after successful AAV-RPE65 therapy (11, 15, 16). In addition, there was no improvement in foveal cone function despite AAV vectors having been delivered to the fovea in the majority of participants (16, 46). These studies highlight the importance of understanding the mechanism of foveal cone degeneration and designing adjunct therapy to slow photoreceptor degeneration in LCA patients in future clinical gene therapy trials. The prominent role of cone opsins in cone degeneration suggests that, in addition to restoring the 11-*cis*-retinal supply, research needs to focus on the folding, trafficking, and degradation properties of cone opsins to design an effective prevention and treatment strategy. Our work demonstrated the importance of M-opsin degradation associated proteasome stress in L/M cone (the dominant cone type in the fovea). We propose to design drugs to help the folding and targeting of M/L-opsins and to enhance proteasome activity to slowdown cone degeneration as an adjunct treatment for gene therapy to achieve the best therapeutic efficacy for LCA (Fig. 4).

## Experimental procedures

### Mice

*M-opsin*^−/−^ mice were generated by CRISPR/Cas9 following a published protocol (47). sgRNA (5’-ATTATCACATTGCTCCCAGG-3’) was designed to target the second exon of mouse M-opsin gene *Opn1mw*. Cas9 plasmid and sgRNA were co-injected into C57Bl/6J mouse embryos at the University of Utah Transgenic/Gene Targeting Core Facility. Mouse tail samples from *Opn1mw*^−/−^ founder mice were prepared and used as PCR template with forward primer (5’-CATAGAGCAAGGAAAAGTGAGGTC-3’) and reverse primer (5’- CCCAGAACGAAGTAGCCATAGAT-3’). The PCR product was gel purified and sequenced to select the founder mice. The pups of the *Opn1mw*^−/−^ founder mice were confirmed by DNA sequencing. *Lrat*^−/−^, *Opn1sw*^−/−^, and *Lrat^−/−^Opn1sw*^−/−^ mice were generated previously (4, 19, 33). *Opn1sw*^−/−^*Opn1mw*^−/−^ mice were produced by crossing the *Opn1sw*^−/−^ and *Opn1mw*^−/−^ lines. *Lrat^−/−^ Opn1sw^−/−^Opn1mw*^−/−^ mice were produced by crossing the *Lrat*^−/−^S-*opsin*^−/−^ and *M-opsin*^−/−^ lines. *Lrat^−/−^Opn1sw^−/−^Ub^G76V^*-*GFP* and *Lrat^−/−^Opn1sw^−/−^Opn1mw^−/−^Ub^G76V^*-*GFP* mice were produced by crossing the *Lrat^−/−^Opn1sw*^−/−^ and *Lrat^−/−^Opn1sw^−/−^Opn1mw*^−/−^ lines, respectively, with the reporter Ub^G76V^-GFP mice. Mice were reared under cyclic light (12 h light/12 h dark).

### Immunohistochemistry

The immunolabeling experiments were performed as previously described (8, 19). Briefly, age-matched mouse eyes were immersion fixed for 2 h using freshly prepared 4% paraformaldehyde in 0.1 M phosphate buffer, pH 7.4, and cryoprotected. Eyes were sectioned at 12-14 μm thickness before being incubated for immunohistochemistry. Primary antibodies were applied to each group of two to four sections in a humidified chamber overnight at 4°C, and were visualized with Alexa 488-, Alexa 647- or Cy3-conjugated secondary antibodies. The sections were viewed using a Zeiss LSM 510 or LSM 800 confocal microscopes. Primary antibodies against S-opsin, M-opsin, cone arrestin, GRK1, and G_α_t2 were described previously (8, 19).

### Western blotting

Retinas from 1-month WT and mutant mice were homogenized by a homogenizer. Protein concentrations were measured by BCA assay (Bio-Rad). 5μg retinal lysates were separated on 10% SDS-PAGE, transferred to a PVDF membrane, and probed with primary antibodies against M-opsin, S-opsin, and Gapdh as described previously (8, 19). The signals were visualized by the ChemiDoc imaging system (Bio-Rad).

### Statistics

All group results are expressed as mean ° SEM. We performed each experiment at least three times and used representative data in the calculations. Comparisons between groups were made using the two-tailed Student’s *t*-test or one-way ANOVA and Tukey’s *post hoc* tests for multiple groups. Levene’s test was used to access homogeneity of variance. Values of ‘N’ were described in figure legends. Statistical significance was denoted with \**P*<0.05, \*\**P*<0.01, \*\*\**P*<0.001 in the figures and figure legends. Statistical analysis was performed with OriginPro.

### Study approval

Animal experiments were approved by the Institutional Animal Care and Use Committees (IACUC) at the University of Utah and Baylor College of Medicine and were in accordance with the Statement of the Association of Research for Vision and Ophthalmology for the Use of Animal in Ophthalmic and Vision Research.

## Acknowledgements

We thank Dr. Wolfgang Baehr for providing the *Lrat*^−/−^ mice and the cone arrestin antibody. Dr. Edward N. Pugh Jr. for providing the *Opn1sw*^−/−^ mice. Dr. Jeannie Chen for providing the S-opsin antibody. This work was supported by NIH grant EY022614 (YF), 2P30EY002520 (Baylor College of Medicine), 5P30EY014800 (University of Utah), the grant from Retinal Research Foundation (YF), the Sarah Campbell Blaffer Endowment in Ophthalmology (YF), and unrestricted grants to the Department of Ophthalmology at the University of Utah and the Department of Ophthalmology at Baylor College of Medicine from Research to Prevent Blindness.

## Author Contributions

H.X. performed most experiments, analyzed the data, and wrote the initial draft. N. P. did sample collections and experimental discussion. X. Z. did some genotyping and mouseline maintenance. O. Z. performed genotyping. Y.F. designed the study, analyzed the data, and wrote the manuscript.

