## Supplementary Figures for "Deletion of M-opsin prevents “M cone” degeneration in a mouse model of Leber congenital amaurosis"

```
atg gcc caa agg ctt aca ggt gaa cag aca ctg gac cac tat gag gat agc acc cat gca
 M   A   Q   R   L   T   G   E   Q   T   L   D   H   Y   E   D   S   T   H   A
agc atc ttc acc tat acc aac agc aac agc acc aaa ggt ccc ttt gaa ggc ccc aat tat
 S   I   F   T   Y   T   N   S   N   S   T   K   G   P   F   E   G   P   N   Y
cac att gct ccc cag gtg ggt gta cca cct cac cag cac ctg gat gat tct tgt ggt cgt
 H   I   A   P   Q   V   G   V   P   P   H   Q   H   L   D   D   S   C   G   R
tgc atc tgt ctt cac taa tgg act tgt gct ggc agc cac cat gag att caa gaa gct gcg
 C   I   C   L   H   -   W   T   C   A   G   S   H   H   E   I   Q   E   A   A
cca tcc act gaa ctg gat tct ggt gaa ctt ggc agt tgc tga cct agc aga gac cat tat
 P   S   T   E   L   D   S   G   E   L   G   S   C   -   P   S   R   D   H   Y
tgc cag cac tat cag tgt tgt gaa cca aat cta tgg cta ctt cgt tct ggg aca ccc tct
 C   Q   H   Y   Q   C   C   E   P   N   L   W   L   L   R   S   G   T   P   S
gtg tgt cat tga agg cta cat tgt ctc att gtg
 V   C   H   -   R   L   H   C   L   I   V
```

**Supplementary Figure 1. The nucleotide sequence of the coding region of M-opsin after CRIPSR mediated insertion.** The 1 bp (C) insertion was shown in red. The first early stop codon due to frame shift was highlighted in yellow.

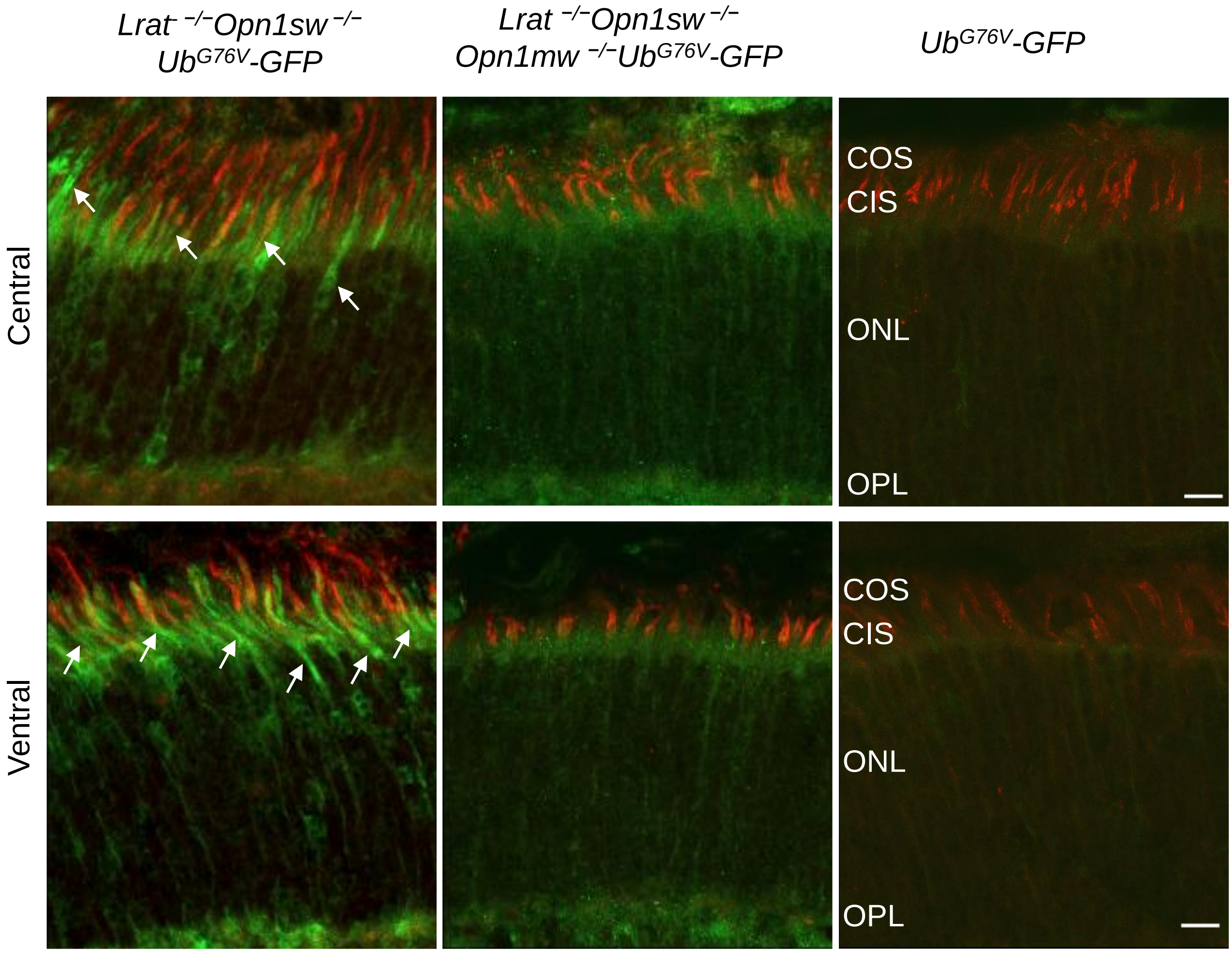


**Supplementary Figure 2. M-opsin deletion reduces proteasomal stress in central and ventral cones of *Lrat* $^{-/-}$*Opn1sw* $^{-/-}$ mice.** The reporter Ub$^{G76V}$-GFP signal from the dorsal retinas of P18 *Lrat* $^{-/-}$*Opn1sw* $^{-/-}$*Ub*$^{G76V}$*-GFP, Lrat* $^{-/-}$*Opn1sw* $^{-/-}$*Opn1mw* $^{-/-}$*Ub*$^{G76V}$*-GFP,* and *Ub*$^{G76V}$*-GFP* control mice were labeled with an anti-GFP antibody (in green). Cones were labeled with rhodamine-PNA (in red). Scale bar, 10 µm.

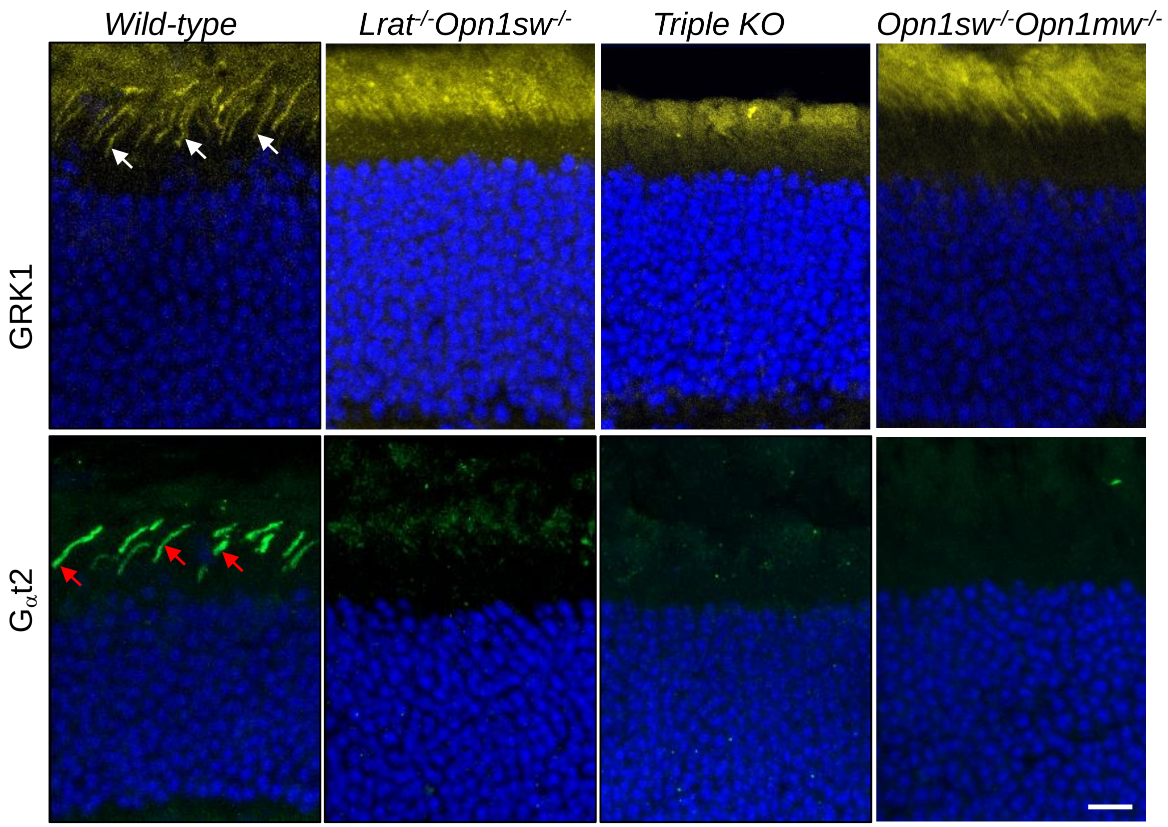


**Supplementary Figure 3. Immunolocalization of cone membrane-associated proteins GRK1 and $G_{\alpha}$t2 in *WT, Lrat$^{-/-}$Opn1sw$^{-/-}$, Lrat$^{-/-}$Opn1sw$^{-/-}$Opn1mw$^{-/-}$, and Opn1sw$^{-/-}$ Opn1mw$^{-/-}$* mice.** 1-month mouse retinal sections were stained with antibodies against GRK1 (in yellow) and $G_{\alpha}$t2 (in green). White arrows and red arrows indicate GRK1 and $G_{\alpha}$t2 signal in WT cones, respectively. Nuclei were stained with DAPI (blue). Scale bar, 10 µm.
